# CD-GPT As a Biological Foundation Model Bridging the Gap between Molecular Sequences Through Central Dogma

**DOI:** 10.1101/2024.06.24.600337

**Authors:** Xiao Zhu, Chenchen Qin, Fang Wang, Fan Yang, Bing He, Yu Zhao, Jianhua Yao

## Abstract

The central dogma serves as a fundamental framework for understanding the flow and expression of genetic information within living organisms, facilitating the connection of diverse biological sequences across molecule types. In this study, we present CD-GPT (Central Dogma Generative Pretrained Transformer), a generative biological foundation model with 1 billion parameters, aiming to capture the sequence relationships between DNA, RNA, and proteins. We model sequences in a unified representational space and employ a shared, multi-molecule vocabulary to narrow their distances in the embedding space effectively. Through extensive pretraining on nucleotide and amino acid sequence data, CD-GPT exhibits exceptional performance in a wide range of predictive and generative downstream tasks, including mono-molecular and multi-molecular analyses. Notably, CD-GPT excels in tasks such as genomic element detection, protein property prediction, RNA-protein interaction identification and also generative tasks like protein generation and reverse translation. The versatility of CD-GPT opens up promising avenues for advanced multi-omics analysis.

## 1 Introduction

The central dogma [1] describes the flow and expression of genetic information within living organisms. Genetic information is first transcribed from DNA to mRNA, then translated into amino acid sequences at the ribosome, and ultimately forms proteins. This coherent process involves the three most crucial molecules in life activities: DNA, RNA, and proteins, which work together to support the diversity and complexity of living organisms.

Advances in sequencing technology have greatly enriched the biological sequence data available to researchers [2, 3], providing support for numerous downstream tasks and research areas, including understanding the regulatory mechanisms of gene expression, identifying genetic variations, and exploring protein modifications. Drawing on the successful experience of pretrained language models [4–6] in the field of Natural Language Processing, which learns the characteristics and patterns of language from vast amounts of text data, biological foundation models [7–9] have also begun to show their potential in addressing related problems. Models based on the Transformer architecture [10] with self-attention mechanism can effectively learn the representations of sequences, bringing more beneficial knowledge transfer to downstream tasks.

DNA, RNA, and proteins each possess a distinct molecular composition, with nucleotides and amino acids exhibiting fundamental chemical differences. Although all can serve as carriers of genetic information, they display significant variability in regulatory mechanisms and functional execution. Additionally, in scientific languages, DNA and RNA are composed of four types of nucleobases, while proteins are made up of twenty different amino acids. These differences lead to distinct embedding representations in foundational models, creating an informative gap that needs to be narrowed.

Current biological foundation models typically focus on pretraining with a single type of molecular sequencing data, such as DNA [11], RNA [12], or proteins [13]. While models like Evo [14] and LucaOne [15] aim to provide a more comprehensive understanding of the entire molecular system within a unified framework, their architectures are often either dominated by one specific molecular type or rely on separate pretraining for different molecular sequences without establishing meaningful connections between them.

As the importance of multi-omics analysis grows, researchers are increasingly focusing on integrating different types of molecular data [16, 17]. However, the limitations of existing models may hinder their ability to fully capture the complex interactions and synergistic effects between various molecular types, ultimately affecting our comprehensive understanding of biological systems. Therefore, developing models capable of truly integrating multi-molecular data has become a key research focus.

In this paper, we introduce CD-GPT, a generative foundation model trained on multi-molecular data (**Fig**.1A-C). CD-GPT employs a unified encoding framework to model data from different molecular types in a comprehensive manner. Compared to mono-molecular models, CD-GPT reduces the distance in the embedding space between various molecular types by leveraging the explicit mapping relationships revealed by the central dogma, creating stronger and more interpretable connections.

**Fig 1.**
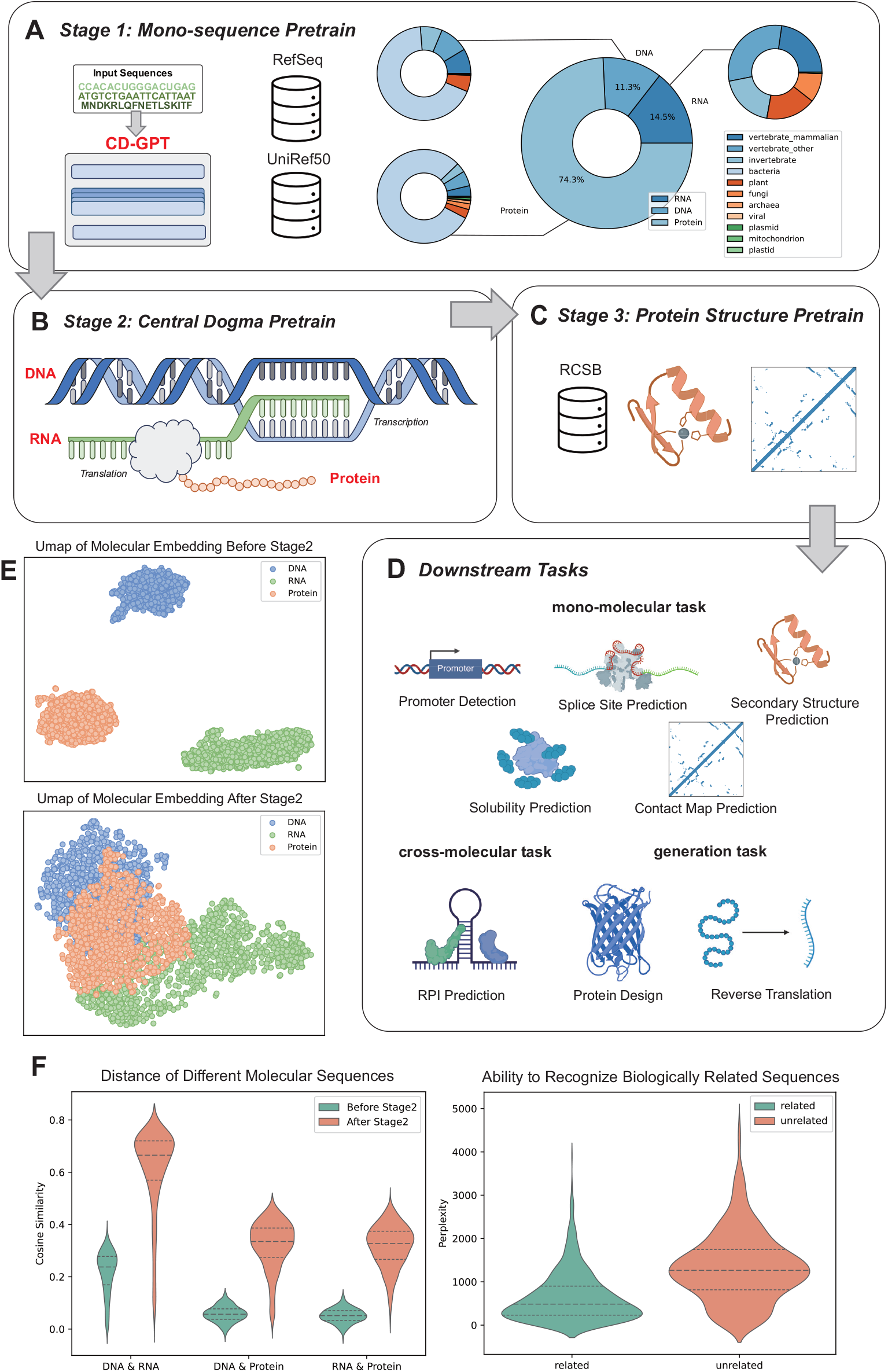
Building a foundation model biological sequences at the full molecular level. CD-GPT underwent three stages of pretrain. **(A)** Firstly, We pretrained CD-GPT on extensive and diverse data set of mono-molecular sequences. **(B)** Subsequently, we pretrained CD-GPT on paired molecular sequences through central dogma, aiming to enable the model to discern the intrinsic connections between various molecular data. **(C)** Lastly, we performed additional pretrain on protein structure data to enhance the model’s deep comprehension of protein secondary structure. **(D)** After these pretrain phases, CD-GPT is capable of handling a variety of molecular and sequence tasks, including mono-molecular tasks, cross-molecular tasks, and sequence generation tasks. **(E)** UMAP visualization showing that the distances between different molecular data in the latent space have been effectively reduced. **(F)** We observed that CD-GPT enables alignment of different molecular types and captures biologically meaningful relationships between these sequences.

We demonstrate that CD-GPT efficiently handles a variety of downstream tasks, including prediction and generation, across the entire molecular level (**Fig**.1D)). First, we show that CD-GPT narrows the gap between sequences from different molecular types at the embedding level, thereby establishing a unified expression framework. Second, CD-GPT achieves outstanding performance on tasks such as promoter detection, splice site prediction, solubility prediction, and RNA-protein interaction prediction. These results indicate that CD-GPT not only learns effective sequence representations during pretraining but also gains a deep understanding of the intrinsic connections between different molecular data types. Finally, in sequence generation tasks such as protein design and reverse translation, CD-GPT demonstrates its potential as a powerful generative model. Through comprehensive evaluation across different molecular and task types, we highlight CD-GPT’s excellent performance and broad application prospects in multi-omics data analysis. This establishes a new paradigm for biological foundation models: modeling at the full molecular level. We hope that CDGPT will contribute to a wider range of biological applications and inspire innovative breakthroughs in life science research.

## 2 Results

### 2.1 CD-GPT narrows the gap between different molecular sequences through central dogma pretrain

In the field of natural language processing, multilingual models effectively address cross-lingual challenges such as machine translation by integrating data from various languages during pretraining [18–21]. Similarly, within intricate biological systems, diverse molecules can be considered as distinct languages. Molecules within organisms engage in information exchange and interaction through a multitude of biological processes. However, due to differences of the representation systems, the embeddings of biological data from different molecular types in the latent space often lie far apart from each other. This observation led us to explore strategies to enrich our model with knowledge from the pretraining phase, thereby bridging the gap between different molecular data.

A unified representational space necessitates a shared foundational vocabulary [20]. Rather than using single-granularity tokenization, which only considers individual nucleotides or amino acids, we draw inspiration from cross-lingual models [22] and develop a more flexible and inclusive system for representing biological sequences. This system utilizes a shared, multi-molecular vocabulary, which facilitates alignment across different molecular data types. Single-granularity tokenization tends to result in sparse spatial distributions that hinder the model’s ability to capture inter-molecular relationships. In contrast, we use Byte Pair Encoding (BPE) [23] to tokenize a mixed molecule corpus, constructing a biological vocabulary that serves as an anchor for aligning diverse sequences through shared symbols.

To integrate the biological principles underlying the central dogma, we designed a specialized dataset during pretraining. This dataset encompasses the flow of genetic information—detailing transcription and translation processes—and pairs related upstream and downstream sequences to enable the model to learn the inherent correlations between molecular sequences.

Our findings, visualized using UMAP, indicate that central dogma pretraining significantly improved the alignment of different molecular sequences in the shared latent space. Prior to pretraining, DNA, RNA, and protein sequences were sparsely distributed, with little to no correlation observed between them. However, after central dogma pretraining, these sequences began to cluster more closely in the embedding space (**Fig**.1**E)**. We further quantified this improvement by calculating the cosine similarity between DNA, RNA, and protein sequences before and after pretraining. After central dogma pretrain, the cosine similarity significantly increased, particularly within triplets of DNA, RNA, and protein sequences (**Fig**.1F left), demonstrating that this pretrain phase effectively reduced the distance between different molecular modalities.

While these observations suggest that central dogma pretraining facilitated the alignment of different molecular types, we also wanted to assess whether the model had captured biologically meaningful relationships between these sequences. To do this, we designed a perplexity-based experiment to evaluate the model’s ability to distinguish between sequences that are biologically related. Perplexity reflects the degree of uncertainty that a model has towards a sequence. For each sequence pair, we concatenated both related and unrelated sequences (e.g., DNA and its corresponding translated protein sequence, versus a randomly selected protein sequence) and calculated their perplexity. A lower perplexity indicates that the model is more confident in its understanding of the connection between the two sequences. Our results show that the model exhibits significantly lower perplexity on sequence pairs that are biologically related (**Fig**.1F right), with a p-value of 2.77 *×* 10^*-*97^. This extremely small p-value indicates a highly statistically significant difference, demonstrating that CD-GPT not only aligned different molecular sequences during pretraining but also learned to recognize meaningful biological relationships between them. The model’s lower perplexity on biologically related pairs further supports the hypothesis that the central dogma pretraining phase helps the model develop biologically informed representations.

### 2.2 CD-GPT is capable of accurately identifying genomic elements

Promoters are specific sequences on DNA that mark the starting point for the transcription of genes into mRNA, representing the first step in the flow of genetic information in the central dogma. Splice donors and acceptors are the exact locations where alternative splicing occurs in the human genome. This mechanism allows for the production of multiple proteins from a single gene through different mRNA splicing variants, enhancing an organism’s adaptability to environmental changes. To evaluate our model’s performance in understanding and predicting these key genetic elements, we employed promoter detection and splice site prediction to test the model’s ability to recognize various genetic elements.

In the tasks of promoter detection and splice site prediction, we followed the dataset division and preprocessing methods of GUE benchmark [11]. We conducted full finetuning on the 1b parameter version of CD-GPT on these specific tasks.

Our model outperformed various existing counterparts on these tasks (**Fig**.2B), including nucleotide transformer [7], DNABERT-2 [11], and Evo [14]. Our results indicate that CD-GPT is not only capable of learning effective sequence representations from a large amount of genomic data but also able to transfer this knowledge to specific downstream tasks, demonstrating excellent generalization ability. Additionally, we observed that the performance of CD-GPT, which underwent central dogma pretrain, was significantly enhanced compared to the version without this process **(Fig.2B**). This finding further confirms the effectiveness of the central dogma pretrain strategy.

**Fig 2.**
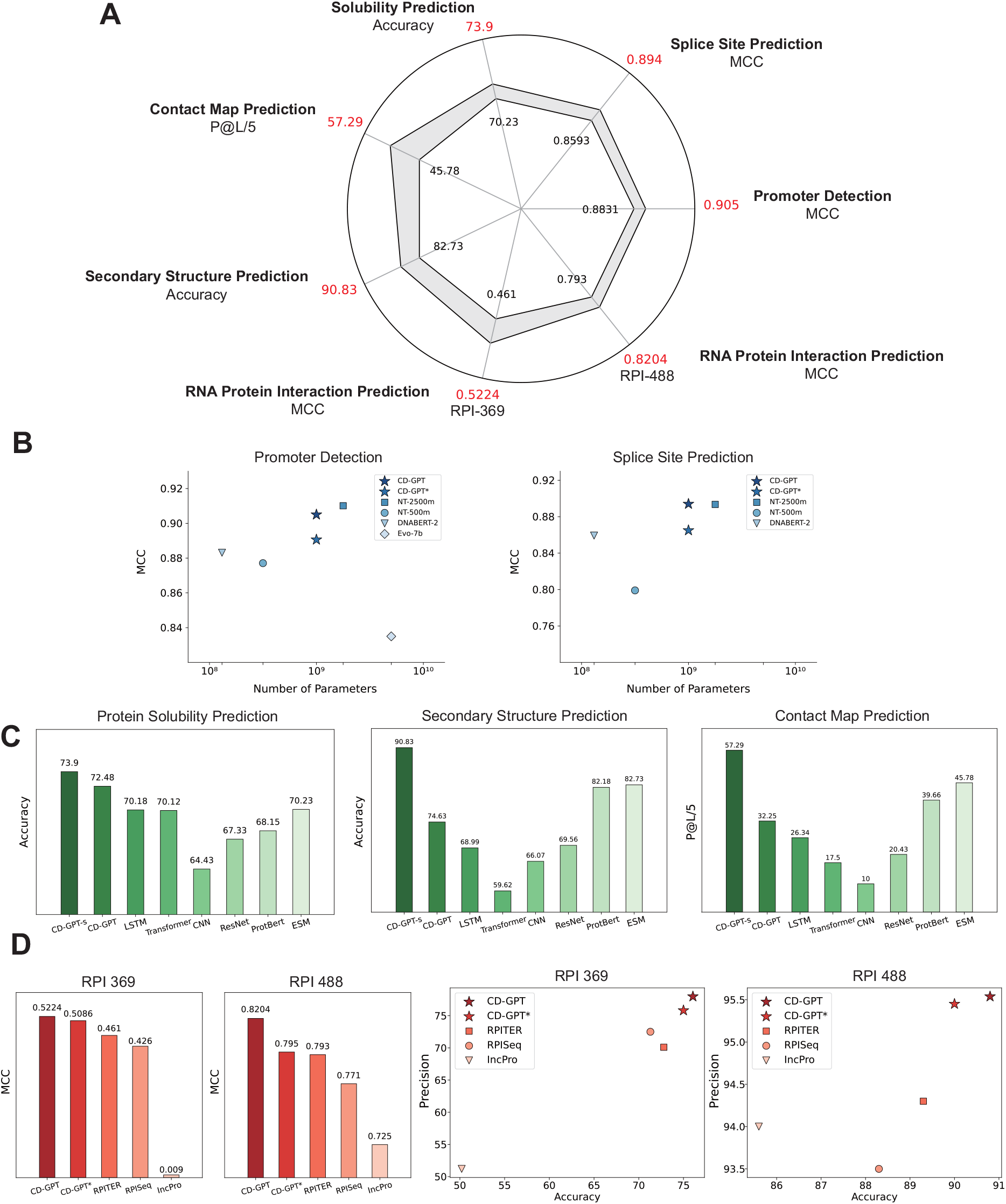
Assessment of CD-GPT’s performance on various downstream tasks. **(A)** CDGPT has shown superior performance over counterparts of similar parameter size in a diverse array of downstream tasks, including promoter detection and splice site prediction **(B)** for genomics data, solubility prediction, contact map prediction and secondary structure prediction **(C)** for protein data, RNA Protein interaction prediction **(D)** for cross molecular data. CD-GPT* refers to a version without central dogma pretrain and CD-GPT-s refers to a version with additional protein structure pretrain.

### 2.3 CD-GPT is able to predict properties and secondary structures of protein

Next, we evaluated CD-GPT’s performance on downstream tasks related to protein data. Proteins are central to all biological functions, as they act as catalysts and execute various cellular processes. Their properties and structures directly influence their biological roles [24]. Therefore, accurately predicting protein properties and structures is crucial for understanding complex cellular mechanisms. To comprehensively assess CD-GPT’s capabilities, we selected several downstream tasks, including solubility prediction, secondary structure prediction, and contact map prediction.

To improve performance on secondary structural prediction tasks, we implemented several design strategies. We customized CD-GPT’s architecture by adding output heads specifically designed for downstream tasks. Additionally, during pretraining, we used a tokenization approach focused on single amino acids at a low ratio, enabling the model to capture finer details at the residue level. For structural predictions, we introduced a phase of pretraining focused on predicting protein secondary structures, helping the model learn key structural patterns essential for downstream tasks (See **Methods** for details).

We then evaluated CD-GPT on solubility prediction, secondary structure prediction, and contact map prediction. Our results show that CD-GPT outperforms other protein-specific models across these tasks. In the solubility prediction task, CD-GPT achieved an accuracy of 73.9%, surpassing the performance of other models. The model also demonstrated exceptional performance in secondary structural prediction tasks, which we attribute to the aforementioned pretraining strategies. When evaluating CD-GPT on solubility prediction, we observed further improvements in predictive performance (**Fig**.2C). This finding reveals that incorporating structural information enables the model to better capture protein characteristics. For example, in solubility prediction, differentiating between hydrophobic and hydrophilic regions is crucial, and the properties of these regions are closely related to the protein’s three-dimensional structure [25]. Consequently, the integration of structural knowledge allows the model not only to recognize sequence features but also to infer indirect characteristics—such as those influencing solubility—that are critical for accurate predictions.

### 2.4 CD-GPT can capture the interactions between different molecular data

Non-coding RNAs (ncRNAs), particularly long non-coding RNAs (lncRNAs), are playing an increasingly critical role in regulating gene expression and participating in cellular signaling pathways [26, 27]. The diverse biological functions of lncRNAs are largely achieved through interactions with specific RNA-binding proteins [28, 29]. Thus, effectively identifying RNA-protein interactions (RPI) is of great significance for deepening our understanding of the biological functions of ncRNAs. By revealing the molecular mechanisms of these interactions, we can better grasp the role of lncRNAs in cellular physiology and pathology, thereby providing new molecular targets and therapeutic strategies for related diseases.

Traditional machine learning methods [30] have certain limitations when dealing with complex biological sequence data, failing to fully capture all useful information in the sequence data. Currently, some methods [31–33] have adopted deep learning technology, achieving better prediction results through networks such as CNNs or LSTMs [34].

However, there is still a relative lack in using large models to handle such tasks. This is because existing large models usually encode based on single-modal data, finding it difficult to process task inputs of multi-modal data. This greatly limits the model’s ability to handle complex biological problems that require the integration of multi-molecular information. Models like Evo [14] attempt to capture system-wide interactions by learning from large genomic regions. This approach is effective to a certain extent, but when confronted with complex biological systems, relying solely on genomic data makes it difficult to fully capture all levels of molecular interactions. Thanks to CD-GPT’s ability to model sequences at the full molecular level, the model can reveal potential connections between different molecular sequences. By concatenating multiple sequences into one, CD-GPT can model data containing multimolecular information directly, as if it were a single sequence. This method not only simplifies the preprocessing but also enhances the model’s ability to capture multimolecular features of complex biological systems.

In comparison with baseline methods, our model has shown strong competitiveness. On the RPI369 and RPI488 [35] datasets, our model has achieved excellent performance (**Fig**.2D). Furthermore, when comparing the performance of two versions of CD-GPT, the model showed significant performance improvement in RPI tasks after central dogma pretrain. This result reveals that by integrating different molecular data, the model can more comprehensively understand the complex relationships between molecules, thereby significantly enhancing the accuracy of predictions.

### 2.5 CD-GPT generates structured and diverse proteins

Protein design aims to create novel proteins with specific biological functions or to optimize and enhance the performance of existing proteins [36]. Designers first craft or refine the amino acid sequence to meet the anticipated functions and structural features of the protein. Subsequently, at the nucleotide level, this amino acid sequence is reversely translated into a DNA sequence, and necessary codon optimization is performed to ensure effective expression of the protein within the target biological system [37].

To evaluate the quality and validity of the generated sequences, we generated 1000 protein sequences and utilized ESM-Fold for structure prediction. We use perplexity and pLDDT as metrics. A lower ppl value indicates that the generated sequence is statistically closer to real data, making it more reasonable. A higher pLDDT value signifies a more orderly structural feature, which is crucial for protein functionality.

We found that the average ppl value of the generated sequences was 8.17, which is close to 6.45 for real sequences. Meanwhile, the average pLDDT value of the generated sequences reached 61.84, with 37.8% of the sequences having a pLDDT value exceeding 70 (**Fig**.3C). These results are also competitive compared to the metrics reported in previous protein generation models [38].

**Fig 3.**
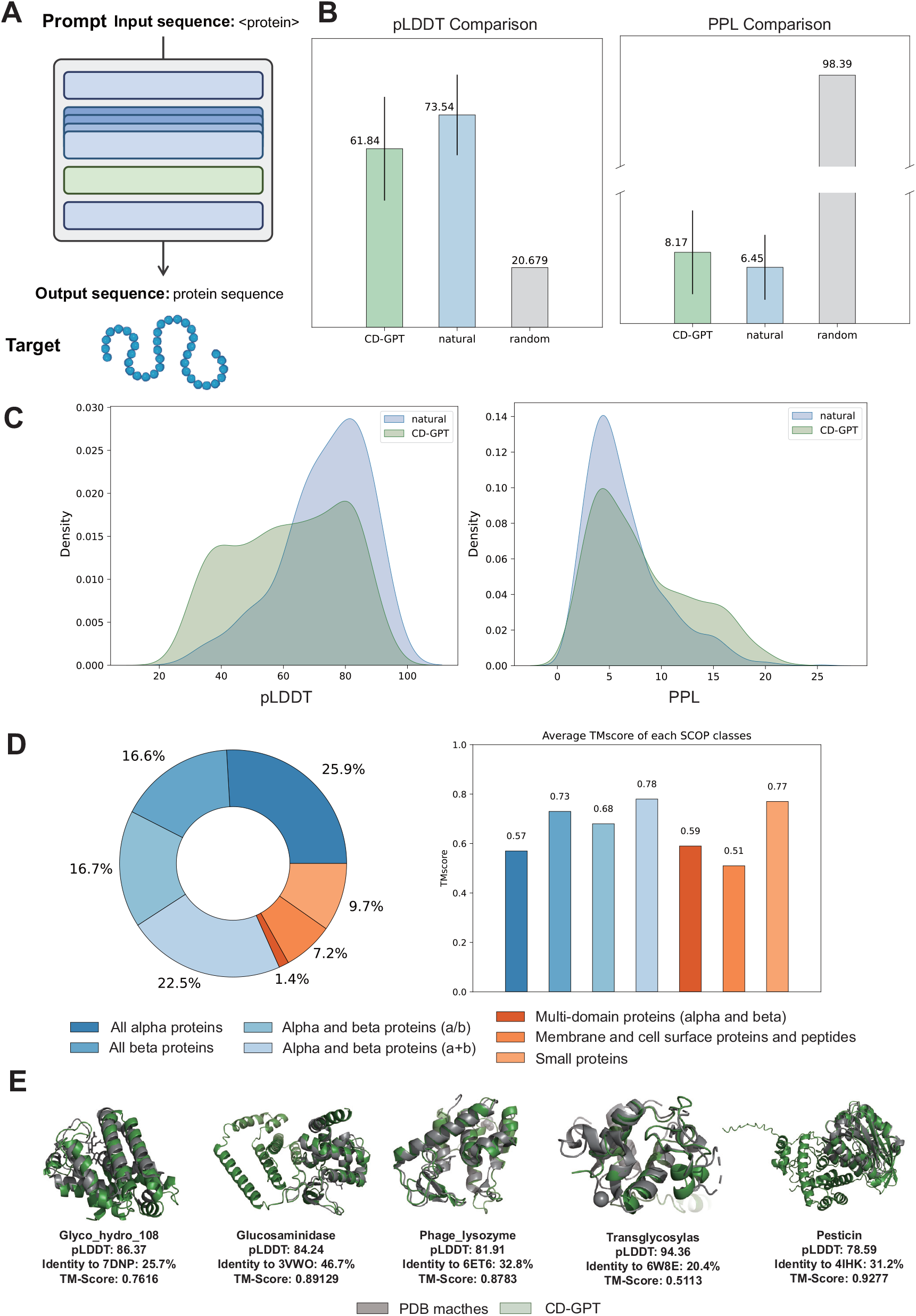
Analysis of protein design results. **(A)** CD-GPT generates corresponding target sequence given prompt label that specifies the protein type. **(B)** Bar charts illustrating the pLDDT and PPL metrics for the CD-GPT generated, natu9ral, and randomly generated proteins. **(C)** Density histogram showing the distribution of the pLDDT and PPL for the CD-GPT generated and natural proteins. **(D)** SCOP classification of generated proteins and average TMscore of each SCOP classes. **(E)** Showcases of generated lysozyme. The structures are predicted by ESM-Fold, and we used Fold-Seek to search structures with high identity.

To further substantiate the structural and functional advantages conferred by the diversity of protein sequences generated by CD-GPT, we generated additional 10000 proteins and employed the FoldSeek to compare the protein structures with those in the SCOP database [39, 40]. Through this process, we identified the target with the highest similarity to the predicted structures as a match. Our analysis revealed that the protein sequences generated by CD-GPT spanned across all SCOP classes (**Fig**.3D), demonstrating the extensive diversity of the generated outcomes. This broad coverage implies that CD-GPT is capable of emulating a variety of protein structures found in nature, offering a wider array of options and possibilities in protein design. To further validate the accuracy of these matches, we calculated the TM-score between the matches. The high TM-score results confirmed the reliability of our structural matches.

We finetuned CD-GPT on the lysozyme family dataset, guiding the model to generate targeted sequences through the use of family member tags. For the generated sequences, we first predicted their structures and identified the highest similarity match in the PDB dataset with FoldSeek. The results show that CD-GPT can successfully generate protein sequences with specific structural features, guided by different prompt tags. The generated sequences exhibit relatively high pLDDT values, suggesting ordered structural features, and display low sequence similarity to known natural proteins (**Fig**.3E).

### 2.6 CD-GPT gains a profound understanding of the central dogma through reverse translation

Translation is a crucial process in the central dogma, marking the key step where genetic information flows from RNA to protein. This process is biologically based on a precise set of rules, where every set of three adjacent nucleotides in messenger RNA, known as a codon, encodes for a specific amino acid. In contrast, reverse translation involves converting a protein sequence back into its corresponding nucleic acid sequence. In the field of bioengineering, it holds significant practical value and importance. Reverse translation can be associated with codon optimization techniques to enhance the expression efficiency of genes within host cells [37, 41].

The complexity of reverse translation lies in the degeneracy of the genetic code, where multiple different codons can encode the same amino acid. This leads to the potential for multiple possible target sequences to be generated during the reverse translation process, increasing the difficulty of accurately deducing the original nucleic acid sequence from the protein sequence. Traditional reverse translation methods often rely on codon usage tables, determining the corresponding codon for an amino acid based on specific codon usage preferences. However, such preferences do not fully reflect the complexity and dynamics within actual biological organisms, such as the abundance of tRNA or the expression level of genes [42]. Ignoring these complex biological factors could result in synthesized gene sequences with low expression efficiency or the production of unintended proteins in the target organism.

We finetuned CD-GPT using translation-related paired sequences from vertebrate mammals, enabling it to follow the prompt instructions for reverse translation tasks and generate corresponding RNA sequences. We evaluated the performance of our model using two metrics: sequence similarity and initiation translation rate. Sequence similarity reflects the model’s learning on codon preferences, while the initiation translation rate is associated with the biological expression efficiency and functionality of the generated sequence [43, 44].

We constructed a codon usage table as a baseline by calculating the frequency of codon occurrences in the finetuning dataset. Upon observing the generation results, we first noted that the model successfully generated valid reverse translation sequences and captured the codon preference (**Fig**.4B), confirming that it has accurately mastered the correspondence between codons and amino acids. In the assessment of sequence similarity, CD-GPT not only surpassed the baseline method but also achieved a high similarity of 84.34% with the natural sequences (**Fig**.4C). This indicates that CD-GPT did not rely solely on codon preferences to generate results but learned a deeper correspondence.

**Fig 4.**
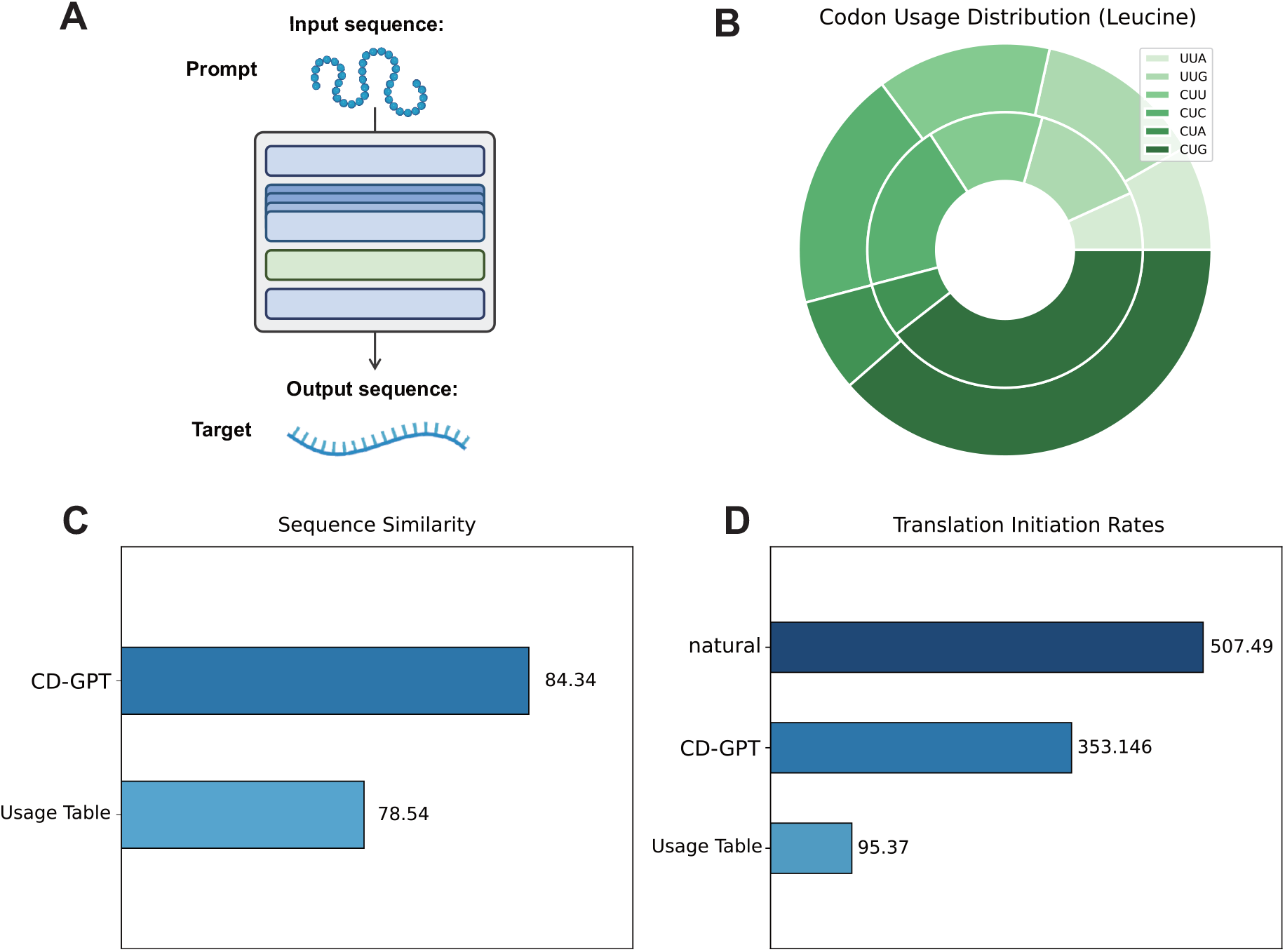
Analysis of reverse translation results. **(A)** CD-GPT generates corresponding target sequences given prompt labels that specify the modality and task type. **(B)** Pie chart illustrating the distribution of codon usage frequencies for leucine in both real sequences and CD-GPT generated sequences. The inner circle represents the natural sequence distribution, while the outer circle depicts the distribution of sequences generated by CD-GPT. **(C)** Bar chart showing the sequence similarity between CD-GPT generated results and the translations based on the codon usage frequency table with the ground truth sequences **(D)** Bar chart showing the translation initiation rates of natural, CD-GPT generated and codon usage table based translation results.

More encouragingly, in the evaluation of the initiation translation rate, CD-GPT not only significantly outperformed the baseline but also approached the measurements of the natural sequences (**Fig**.4D). This finding reveals that the sequences generated by our model have higher potential and functionality in biological expression, which is crucial as it suggests that the generated sequences are more likely to achieve accurate expression and fulfill the desired biological functions within biological organisms.

## 3 Discussion

In this paper, we introduce CD-GPT, a generative foundation model pretrained on comprehensive data at the full molecular level. Through a unified encoding framework, CD-GPT is capable of modeling various types of molecular data in an integrated manner. To support the rationality of this approach, we demonstrate its interpretability by measuring the distance between molecular sequences in the latent space, showcasing how the model captures relationships across diverse molecular modalities.

CD-GPT’s performance across a variety of downstream tasks demonstrates its robust ability to predict molecular properties. By learning cross-modal representations during the pretraining phase, CD-GPT effectively predicts the interactions and properties of different molecular types. This multi-modal framework empowers CD-GPT to address complex, multi-molecular tasks and positions it as a valuable tool for advancing multi-omics research. These results emphasize the necessity of integrating diverse data sources to fully understand the intricate interactions within biological systems and gain deeper biological insights.

Despite the promising results, CD-GPT, as the first generative foundation model for biological data at the full molecular level, still has significant room for improvement, and its potential can be explored further in several directions.

First, according to the scaling laws of generative models [45], the performance of such models is positively correlated with the scale of both the model’s parameters and the data used. Currently, CD-GPT operates at a scale of 1 billion parameters, suggesting that scaling up the model size and incorporating more diverse training data could lead to substantial performance improvements.

Second, while we focused on the RNA-to-protein translation process in the context of the central dogma, the central dogma encompasses broader molecular interactions.

For example, RNA splicing, a process that removes introns and joins exons to form mature mRNA [46], plays a critical role in gene expression regulation and is involved in various biological functions and diseases [47]. Additionally, in retroviruses, RNA can be reverse-transcribed into DNA, a process catalyzed by reverse transcriptase [48], which is crucial for viral replication. Incorporating these complex processes into CD-GPT’s learning framework could further enhance its ability to model molecular dynamics and provide insights into viral mechanisms and gene regulation.

The current model primarily learns relationships between sequences at the RNA, DNA, and protein levels, but does not sufficiently capture interactions at lower levels, such as atom-level or residue-level interactions. These finer-scale interactions are crucial for understanding protein folding, molecular docking, and other critical biological processes. As a result, CD-GPT may struggle to predict precise molecular behaviors or to generalize when simulating interactions at these smaller scales, which is a common challenge faced by foundation models in biology.

Moreover, the complexity of biological regulatory mechanisms, which are inherently dynamic and context-dependent, presents another significant challenge. While CDGPT excels at capturing broad molecular relationships, its ability to simulate and predict regulatory dynamics, such as gene expression and protein modification, remains limited by the specificity and diversity of the training data. The lack of comprehensive datasets that encompass the full range of regulatory processes makes it difficult for the model to fully capture the subtleties of regulatory interactions in real biological environments.

To address these limitations, we plan to enhance CD-GPT’s learning framework by incorporating more detailed molecular interactions at the atomic and residue levels. Additionally, future iterations of CD-GPT will aim to integrate more complex biological processes to build a more accurate model of multi-molecular interactions. Projects like The Cancer Genome Atlas [49] and the Human Cell Atlas [50] offer a wealth of data that can enrich our model and enable more context-aware predictions of gene regulatory mechanisms. These efforts will pave the way for the development of more sophisticated, context-sensitive models that can provide deeper insights into the dynamics of biological systems.

In summary, with ongoing improvements in data integration and model refinement, we anticipate that CD-GPT and similar models will offer new perspectives for understanding the complexity of biology and will serve as powerful tools for researchers exploring the vast potential.

## 4 Methods

### 4.1 Model details

#### 4.1.1 tokenization

We constructed our vocabulary using the Byte Pair Encoding (BPE) algorithm [23], which incrementally merges frequent pairs to build up the vocabulary over iterations. Retaining special symbols such as digits and letters on top of this BPE vocabulary allows for subsequent finetuning and sequence generation tasks. This approach culminates in a comprehensive vocabulary of 64,000 tokens with length ranging from 1 to 16.

We specifically randomly sampled DNA, RNA, and protein sequences from various model organisms, including Homo sapiens, Mus musculus, Danio rerio, Drosophila melanogaster, Caenorhabditis elegans, Saccharomyces cerevisiae, Escherichia coli, and Arabidopsis thaliana. The sampling was done in a 5:3:1 ratio, reflecting the proportionate lengths of the DNA, RNA, and protein sequences, respectively. We constructed a vocabulary using SentencePiece [51] and manually added special symbols such as numbers and letters to enhance the universality and applicability of the vocabulary.

#### 4.1.2 architecture

CD-GPT adopted a transformer decoder-only architecture [10], which consists of three main components: an embedding layer, a stack of transformer decoder layers, and an output head (**Fig**.5A).

**Fig 5.**
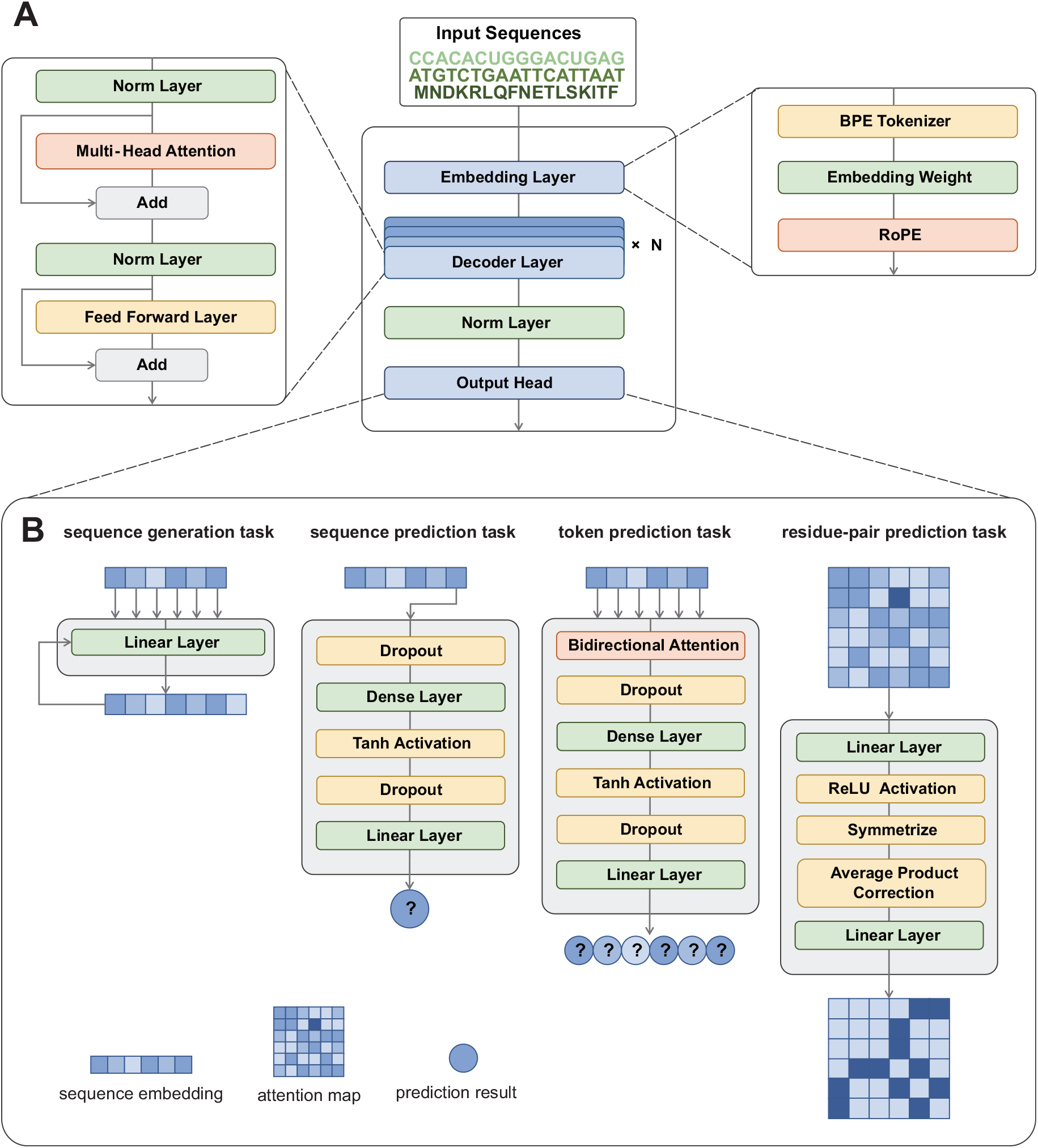
Detailed architecture of CD-GPT. **(A)** CD-GPT consists of three main components: an embedding layer, a stack of transformer decoder layers, and an output head. **(B)** According to different types of downstream tasks, We apply different output heads to get the generation or prediction results.

In the embedding layer, the input sequence, represented as a list of tokens by the BPE tokenizer, is transformed into a learnable vector representation. This vector representation is then fed into a series of decoder layers to obtain a richer semantic representation. Each decoder layer is composed of a multi-head causal attention layer and a feed-forward layer. The self-attention mechanism [10] is the core component of this layer and can be formulated as:

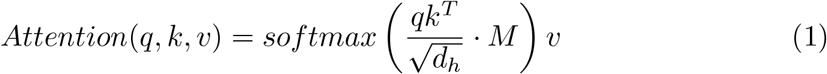

where *q, k, v* denote query, key, value representation and *d*_*h*_ represents the hidden size of multi-head attention. We used a lower triangular mask *M* during the attention calculation, which sets the dot product results of the key and query for future positions to a very large negative number before the attention score calculation, resulting in almost zero after the softmax function.

We employ rotary positional encoding [52] to provide the model with positional information for each token in the sequence. It effectively conveys positional information within the model by representing each position’s encoding as a rotation operation on the complex plane.

After processing by the embedding and decoder layers, the input sequence is transformed into a vector embedding that carries functional and structural information. Depending on the downstream task, we then apply the corresponding output head to obtain the results (**Fig**.5B).

For **sequence generation** tasks, we use a prediction head that maps each token in the sequence to the probability distribution of its subsequent token. The output head includes a linear layer with an input dimension of the model’s embedding dimension and an output dimension equal to the length of the vocabulary. When generating a sequence, we take the output result of the last token and sample from it to produce the generated token. We combine the generated token with the original sequence to form a new sequence and continue sequence generation by repeating the above process.

For **sequence prediction** tasks, we use a prediction head that maps the last token of the sequence to the probability distribution of the category. The output head includes a linear layer with an input dimension of the model’s embedding dimension and an output dimension equal to the number of predicted categories. During prediction, we input the representation of the last token into the linear layer to obtain the prediction result, as the representation of the last token has already exchanged information with the entire sequence in the previous attention computation process.

For **token prediction** tasks, we use a prediction head that maps each token in the sequence to the probability distribution of the category. The output head includes a bidirectional attention layer and a linear layer. We have added a bidirectional attention layer in front of the prediction head, allowing tokens earlier in the sequence to exchange information with later tokens, thereby achieving improved predictive results.

For **residue-pair prediction** tasks, we take the attention matrix as input and use a prediction head to obtain the probability distribution of the category to which each residue pair belongs. The output head includes a two-layer MLP that maps the high-dimensional attention map to the two-dimensional predicted categories. We also use symmetrization and average product correction operations to symmetrize and normalize the attention map [53].

### 4.2 Dataset details

#### 4.2.1 pretraining dataset

The pretraining data corpus for CD-GPT is primarily sourced from the RefSeq [54] and UniRef50 [55]. We initially acquired a comprehensive dataset that covers 14 different organisms, originally provided in FASTA and GenBank file formats. The FASTA files contain the sequence information, while the GenBank files contain detailed annotations of the sequences.

In the first phase of pretraining, we utilized the mono-sequences for training. We removed sequences containing invalid symbols like ‘N’, and then we performed character-level tokenization on the sequences with a probability of 5%. Finally, we obtained 39,771,575 DNA sequences, 51,237,428 RNA sequences, and 26,238,387 protein sequences for pretraining.

In the second phase of pretraining, we employed paired sequences that involve the translation process of the central dogma. Based on the sequence annotations in the GenBank files of RNA data, we pinpointed the CDS regions and forward strands of their upstream DNA sequences, pairing them with the corresponding translated protein sequences to form a complete set of paired sequences. Utilizing this method, we acquired a total of 337,780,146 high-quality sequence pairs, providing a rich corpus for the second phase of pretraining.

We performed additional training on a vast array of PDB data from RCSB [56], thereby integrating structural information into CD-GPT. Utilizing the spatial structures of proteins recorded in PDB files, we calculated their secondary structures and contact maps, ultimately compiling a dataset of 595,108 proteins.

#### 4.2.2 evaluation dataset

For the promoter detection and splice site prediction tasks, we utilized the datasets and division from the GUE benchmark [11].

##### Promoter Detection

Promoter detection dataset is downloaded from the Eukaryotic Promoter Database [57]. Sequences are extracted to span from 249 base pairs upstream to 50 base pairs downstream of the transcription start site. These sequences encompass both TATA-containing and TATA-less promoters. and is extracted to be -249 +50 bp around the transcription start site from both TATA and non-TATA promoters. These two datasets are merged to form the final dataset as promoter class. Non-promoter class dataset is constructed by randomly selecting sequences outside of promoter regions but with TATA motif or randomly substituted sequences without TATA motif. All the datasets above are merged to obtain a final dataset.

##### Splice Site Prediction

For the splice site prediction task, the dataset is sourced from SpliceFinder [58] and includes 400 bp long sequences extracted from the Ensembl GRCh38 human reference genome. To increase the complexity of this task, the dataset is reconstructed by iteratively adding adversarial examples.

For the protein solubility prediction, secondary structure prediction, and contact map prediction tasks, we employed the datasets and division from the PEER benchmark [59].

##### Solubility Prediction

The solubility prediction dataset originates from PROSO II [60] and DeepSol [61]. The dataset consists of 58689 soluble and 70954 insoluble protein sequences. CD-HIT [62, 63] is first used to decrease sequence redundancy with a maximum sequence identity of 90%. Then protein sequences with a sequence identity of *≥* 30% to any sequence in the test set are excluded from the training set to form the final training dataset. We follow the criteria established in the pepcDB database used in the PROSO II for our experiment setup. For each construct, experimental results and status history are recorded in pepcDB. All constructs that achieved the Soluble status or subsequent stages are considered soluble(Soluble, Purified, Crystallized, heteronuclear single quantum coherence, Structure, and In PDB).

##### Secondary Structure Prediction

The training set for the secondary structure prediction task comes from NetSurfP2.0 [64], which is filtered such that no two proteins have greater than 25% sequence identity. The test set is derived from CB513 [65]. It is filtered at 25% sequence identity against the training set to assess the generalization across dissimilar protein sequences.

##### Contact Map Prediction

For the contact map prediction task, we use ProteinNet dataset for training and CASP12 test set for evaluation. The test set is filtered against the training set at a 30% sequence identity. We report precision of the L/5 most likely contacts for long-range contacts as metric.

For the RNA protein interaction prediction task, we used the widely recognized benchmark datasets RPI369 and RPI488 [35] from previous studies.

##### RNA Protein Interaction Prediction

These datasets are extracted from the RNA-protein interaction complexes in the RNA-protein interaction database PRIDB or PDB. If there is a protein atom and an RNA atom with the distance between them being less than the specified distance threshold, then the protein and RNA form an interaction pair. Additionally, RPI488 consists of lncRNA-protein interaction pairs, where the RNAs are ncRNAs longer than 200 nt. We followed previous study using 5-fold cross-validation and reported the average performance.

##### Unconditional Protein Generation

For the unconditional protein generation task, we reserved a part of our pretrain protein dataset, ensuring it was not used during the training phase. We split this reserved dataset into two parts: 90% was used for finetuning our model, and the remaining 10% was used as a natural control set. During fine-tuning, we added a ***<*protein*>*** tag before each sequence in the fine-tuning set. This prompt allows the model to recognize that it should generate a protein sequence when given the ***<*protein*>*** tag during inference. We used FoldSeek to compare the protein structures with SCOP 2.08 filtered at 95% dataset.

##### Lysozyme Generation

For the lysozyme generation task, we leveraged a dataset sourced from Progen [68]. This dataset encompasses five protein families derived from the Pfam [69] database, which includes phage lysozyme, pesticin, glucosaminidase, the glycoside hydrolase family 108, and transglycosylase.

##### Reverse Translation

For the reverse translation task, we constructed a finetuning dataset using mammalian sequencing data from the RefSeq, which includes 11,598,787 protein-pre-mRNA reverse translation sequence pairs, and we set aside 100 pairs for testing. Each pair consists of a protein and its corresponding pre-mRNA, with the mRNA containing only the coding sequence region.

### 4.3 Training details

#### 4.3.1 pretraining objectives

We employed next token prediction [70] as our pretraining task, where the model predicts the subsequent token based on the preceding tokens. We utilized cross-entropy as the loss function to optimize the model parameters, which can be formulated as:

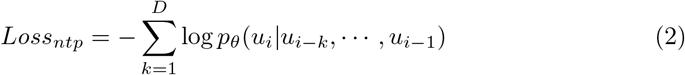

where *u*_*i*_ represents the *i*-th token in the given sequence.

**Table 1.**
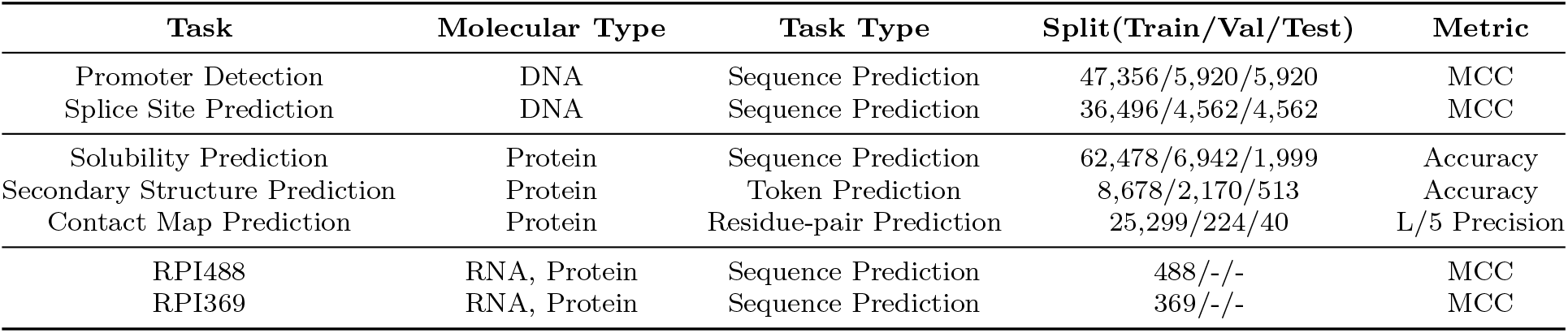
Statistics of the evaluation datasets. We show each task with its related molecular type, task type, the size of each split and metric used for evaluation.

We pretrained CD-GPT with a two-phase process to enhance its performance step by step. These two phases are named **Mono-sequence Pretrain** and **Central Dogma Pretrain**.

In the **Mono-sequence Pretrain**, CD-GPT was trained directly on the mono-sequences of different molecular types. During this phase, the sequences had a 5% chance of being tokenized at individual nucleotides or amino acids level.

In the **Central Dogma Pretrain**, we employed the idea of the central dogma by pairing RNA sequences with protein sequences. We selected sequence pairs from the training data that involved the translation process and organized them into a coherent training corpus.

We conducted additional protein structure pretrain on a vast amount of PDB data. We adopted a multi-task training strategy and trained CD-GPT to simultaneously tackle next token prediction, contact map prediction, and secondary structure prediction tasks. The loss function can be formulated as:

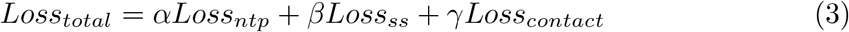

where *α, β, γ* are set to 0.1, 0.8, 0.1 respectively during training.

#### 4.3.2 pretraining settings

We trained CD-GPT on 8 NVIDIA A100 40G GPUs. CD-GPT consumed 206.095 billion tokens during the first phase of pre-training, which lasted for 20 days and consumed 139.274 billion tokens during the second phase of pre-training, which lasted for 14 days.

We employed data parallelism during both phases of pretraining to promote the efficiency and stability of the training process. Additionally, we leveraged the FusedAdam optimizer, which incorporates weight decay to prevent overfitting, along with a learning rate scheduling strategy to dynamically adjust the learning rate throughout the training epochs.

#### 4.3.3 finetuning objectives

In sequence prediction tasks, the model is required to predict one of several possible labels for an input sequence. Tasks such as promoter detection, splice site prediction, solubility prediction, and RNA-protein interaction prediction can be classified under this category. The finetuning loss can be formulated as:

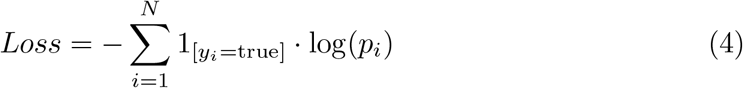

where *N* is the total number of labels, *y*_*i*_ is an indicator that is 1 if label *i* is the correct classification for the input sequence, and *p*_*i*_ is the predicted probability that the input sequence belongs to label *i*.

In token prediction tasks, the model needs to predict one of several possible labels for each token in the input sequence. Secondary structure prediction can be classified under this category. The finetuning loss can be formulated as:

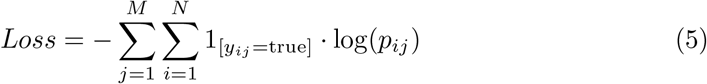

where *M* is the total number of tokens in the sequence, *N* is the total number of labels, *p*_*ij*_ is the predicted probability that *j*-th token belongs to label *i*.

In residue pair prediction tasks, the model needs to predict one of several possible labels for each residue pair of the input sequence. Contact map prediction can be classified under this category. The finetuning loss can be formulated as:

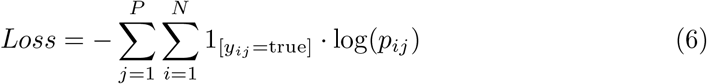

where *P* is the total number of residue pairs in the input sequence. *N* is the total number of labels, *p*_*ij*_ is the predicted probability that *j*-th residue pair belongs to label *i*.

#### 4.3.4 prompt details

We define a “prompt” as the input sentence during training and inference. The prompts vary across different pretraining phases or downstream tasks. Here is a detailed explanation:

In the **Mono-sequence Pretrain**, the prompt are directly the mono-sequences:

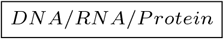

In the **Central Dogma Pretrain**, the prompt is designed as:

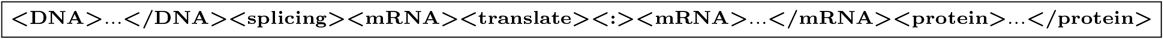

Or

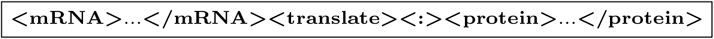

where … represents the corresponding molecular sequences.

In the training phase of **Unconditional Protein Generation** task, the prompt is designed as:

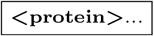

where … represents protein sequences.

In the inference phase of **Unconditional Protein Generation** task, the prompt is only left with a protein tag:

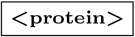

In the training phase of **Lysozyme Generation** task, the prompt is designed as:

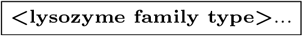

where lysozyme family type can be **Glucosaminidase, Glyco hydro 108, Pesticin, Phage lysozyme** and **Transglycosylas**.

In the inference phase of **Lysozyme Generation** task, the prompt is only left with a lysozyme tag:

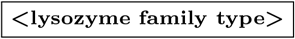

In the training phase of **Reverse Translation** task, the prompt is designed as:

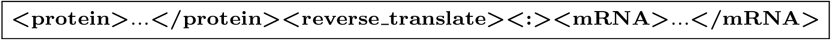

where … represents protein sequences.

In the inference phase of **Reverse Translation** task, the prompt is truncated as:

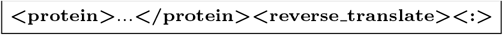

### 4.4 Benchmark details

For most of comparative methods discussed in this paper, due to the use of an identical downstream task dataset partitioning, we directly cited the best performance reported in the original literature.

In the evaluation of Evo [14], we utilized its official implementation on https://github.com/evo-design/evo and its released checkpoints. Referring to the approach taken in this paper for sequence prediction tasks, we selected the representation of the last token in the sequence as the input for the prediction head. We tested Evo on promoter detection task and splice site prediction task. During evaluation, we found Evo difficult to converge on the splice site prediction task, so we only reported its performance on promoter detection task.

In the evaluation of reverse translation task, we employed OSTIR [71] to calculate the Translation Initiation Rates for RNA synthesis. OSTIR leverages the ViennaRNA Package for conducting the requisite free energy calculations, and its entire workflow is freely accessible as open source.

### 4.5 Code availability

The codebase and model checkpoint for CD-GPT is publicly available at Github (https://github.com/TencentAI4S/CD-GPT) and Zenodo [72] (https://doi.org/10.5281/zenodo.14099687).

### 4.6 Data availability

All data used in this study are publicly available and the details are fully illustrated in the Methods. The promoter detection dataset and splice site prediction dataset were downloaded from https://github.com/MAGICS-LAB/DNABERT_2. The solubility prediction dataset was downloaded from https://miladeepgraphlearningproteindata.s3.us-east-2.amazonaws.com/peerdata/solubility.tar.gz. The sec-ondary structure prediction dataset was downloaded from http://s3.amazonaws.com/songlabdata/proteindata/data_pytorch/secondary_structure.tar.gz. The contact map prediction dataset was downloaded from https://miladeepgraphlearningproteindata.s3.us-east-2.amazonaws.com/data/proteinnet.tar.gz. The RPI prediction datasets were downloaded from https://github.com/Pengeace/RPITER.

