## Supplementary File for "CD-GPT As a Biological Foundation Model Bridging the Gap between Molecular Sequences Through Central Dogma"

### Supplementary information

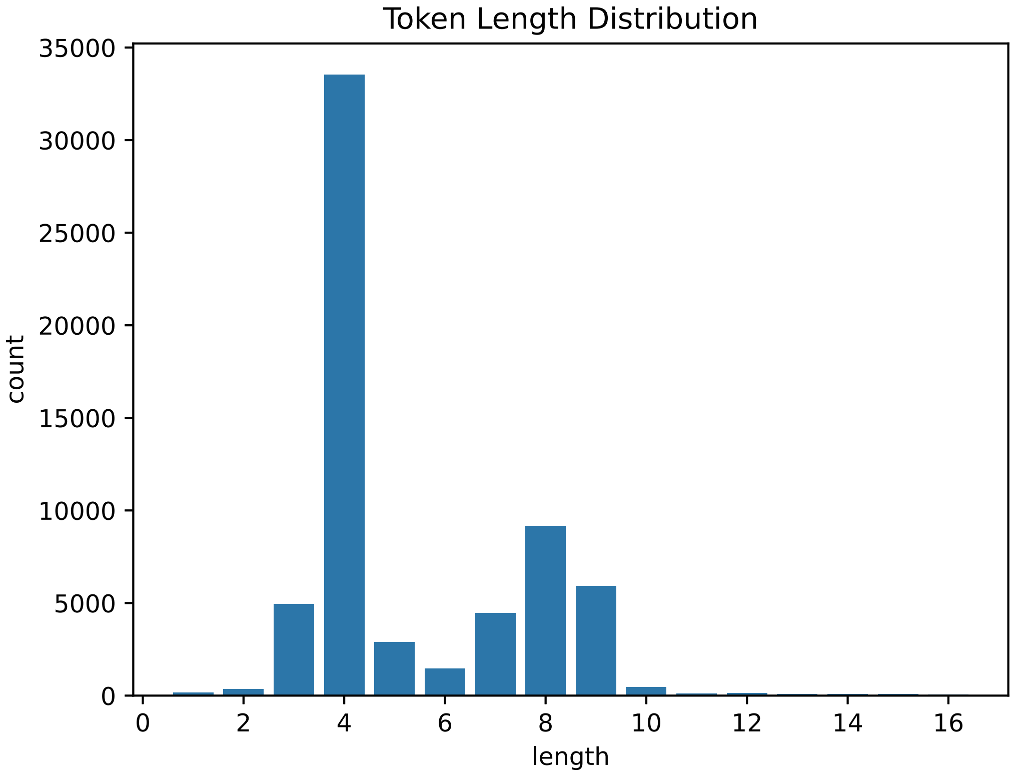


**Supplementary Figure 1: Histogram of token length in the vocabulary.** We constructed vocabulary with variable token length using the BPE algorithm.

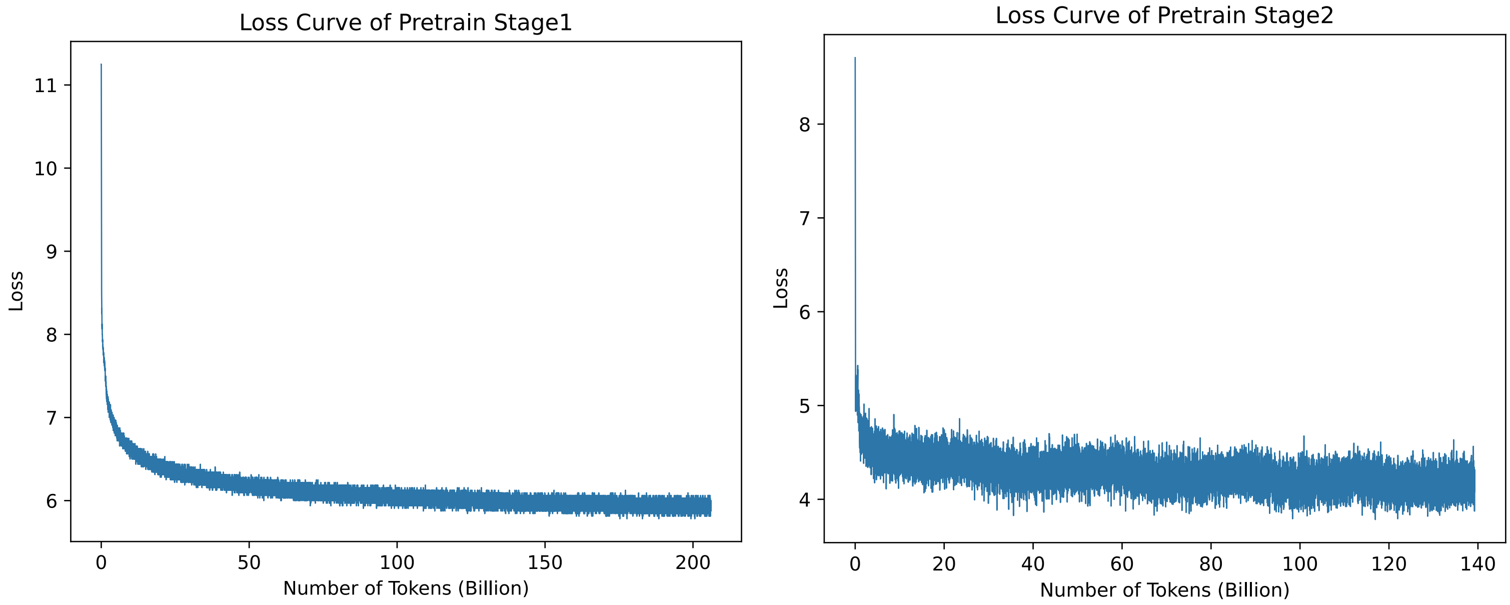


**Supplementary Figure 2: Training loss curves during different pretrain phases.** CD-GPT exhibited a steady decline in loss during both the pretrain phases, indicating effective learning and convergence.

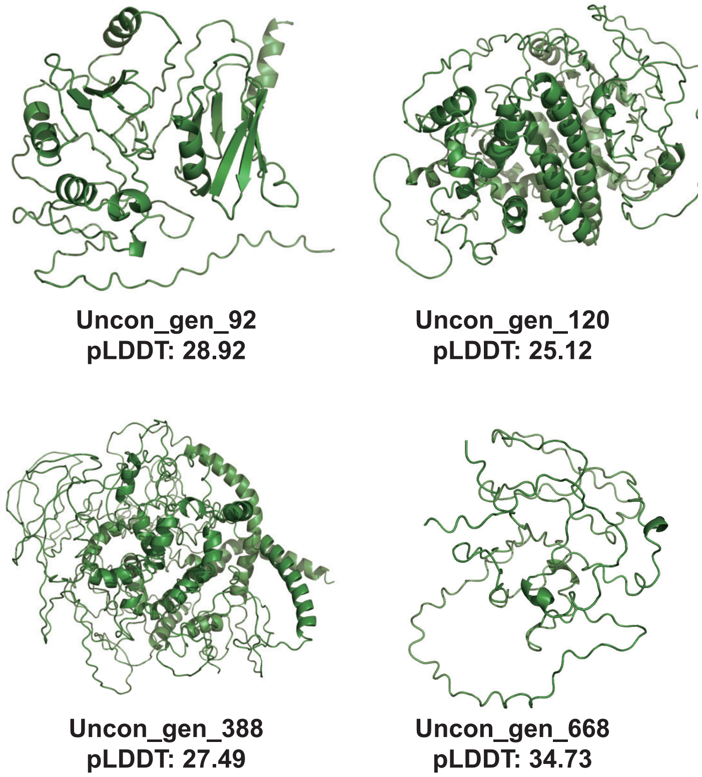


**Supplementary Figure 3: Examples of intrinsically disordered proteins generated by CD-GPT.**

**Supplementary Table 1:** Hyperparameters settings for pretrain and architecture of CD-GPT.

| **Stage1: Mono-sequence Pretrain** | |
|---|---|
| sequence length | 1024 |
| batch size | 40 |
| data parallel | 8 |
| learning rate | 3.00E-05 |
| warmup iter | 10000 |
| weight decay | 0.01 |
| optimizer | FusedAdam |
| optimizer betas | 0.9, 0.999 |
| scheduler | WarmupCosineLR |
| checkpoint activation | TRUE |
| num layers | 12 |
| num heads | 24 |
| embedding dim | 2304 |
| vocab size | 64000 |
| params | 1024M |
| **Stage2: Central Dogma Pretrain** | |
| learning rate | 1.00E-05 |
| batch size | 32 |

**Supplementary Table 2:** Statistics of the pretraining dataset.

| **Pretrain Stage** | **Data Type** | **Amount** | **Data Source** |
|---|---|---|---|
| Mono-sequence Pretrain | DNA | 39,758,549 | RefSeq |
| | RNA | 51,059,612 | |
| | Protein | 262,266,016 | UniRef50 |
| | **Total** | **353,084,177** | |
| Central Dogma Pretrain | DNA-Protein pair | 293,571,497 | RefSeq |
| | mRNA-Protein pair | 44,208,649 | |
| | **Total** | **337,780,146** | |
| Protein Structure Pretrain | Protein | 595,108 | RCSB |

**Supplementary Table 3: Performance comparison of different models on DNA and protein tasks.** CD-GPT* refers to a version with pretrain Stage1. CD-GPT refers to a version with pretrain Stage1&2. CD-GPT-s refers to a version with pretrain Stage 1&2&3. The best and second best results are shown in **bold** and <u>underlined</u> respectively.

| **Task** | **Model** | **Metric** | **Performance** |
|---|---|---|---|
| Promoter Detection | CD-GPT | MCC | <u>0.905</u> |
| | CD-GPT* | | 0.8906 |
| | NT-2500m [1] | | **0.9101** |
| | NT-500m | | 0.8771 |
| | DNABERT-2 [2] | | 0.8831 |
| | Evo [3] | | 0.835 |
| Splice Site Prediction | CD-GPT | MCC | **0.894** |
| | CD-GPT* | | 0.865 |
| | NT-2500m | | <u>0.8935</u> |
| | NT-500m | | 0.7971 |
| | DNABERT-2 | | 0.8593 |
| Solubility Prediction | CD-GPT-s | Accuracy | **73.9** |
| | CD-GPT | | <u>72.48</u> |
| | LSTM | | 70.18 |
| | Transformer | | 70.12 |
| | CNN | | 64.43 |
| | ResNet | | 67.33 |
| | ProtBert [4] | | 68.15 |
| | ESM [5] | | 70.23 |
| Secondary Structure Prediction | CD-GPT-s | Accuracy | **90.83** |
| | CD-GPT | | 74.63 |
| | LSTM | | 68.99 |
| | Transformer | | 59.62 |
| | CNN | | 66.07 |
| | ResNet | | 69.56 |
| | ProtBert | | 82.18 |
| | ESM | | <u>82.73</u> |
| Contact Map Prediction | CD-GPT-s | L/5 Precision | **57.29** |
| | CD-GPT | | 32.25 |
| | LSTM | | 26.34 |
| | Transformer | | 17.5 |
| | CNN | | 10 |
| | ResNet | | 20.43 |
| | ProtBert | | 39.66 |
| | ESM | | <u>45.78</u> |

**Supplementary Table 4: Performance comparison of different models on RPI369 and RPI488.** CD-GPT* refers to a version with pretrain Stage1. CD-GPT refers to a version with pretrain Stage1&2. The best and second best results are shown in **bold** and <u>underlined</u> respectively.

| Task | Model | Performance | | |
|---|---|---|---|---|
| | | MCC | Accuracy | Precision |
| RPI369 | CD-GPT | **0.5224** | **76.054** | <u>77.976</u> |
| | CD-GPT* | <u>0.5086</u> | <u>75</u> | 75.814 |
| | RPITER [6] | 0.461 | 72.8 | 70.1 |
| | RPISeq [7] | 0.426 | 71.3 | 72.5 |
| | IncPro [8] | 0.009 | 50.2 | 51.2 |
| | IPMiner [9] | 0.428 | 70 | **84** |
| RPI488 | CD-GPT | **0.8204** | **90.8** | **95.538** |
| | CD-GPT* | <u>0.795</u> | <u>90</u> | <u>95.45</u> |
| | RPITER | 0.793 | 89.3 | 94.3 |
| | RPISeq | 0.771 | 88.3 | 93.5 |
| | IncPro | 0.725 | 85.6 | 94 |
| | IPMiner | 0.793 | 89.3 | 95.1 |

**Supplementary Table 5: Results of ablation studies on different pretrain phases.** For DNA and RPI tasks, we assessed the efficacy of central dogma pretrain (Stage2). For protein tasks, we assessed the efficacy of protein structure pretrain (Stage3).

| Model CD-GPT | Promoter MCC | Splice Site MCC | Solubility Acc | Secondary Structure Acc | Contact Map P@L/5 | RPI369 MCC | RPI488 MCC |
|---|---|---|---|---|---|---|---|
| w/ Stage1 | 0.8906 | 0.865 | - | - | - | 0.5086 | 0.795 |
| w/ Stage1&2 | 0.905 | 0.894 | 72.48 | 74.63 | 32.25 | 0.5224 | 0.8204 |
| w/ Stage1&2&3 | - | - | 73.9 | 90.83 | 57.29 | - | - |

**Supplementary Table 6:** Qualitative comparison of different sequence generation models. *Evo generates proteins at an indirect manner by generating coding sequences.

| Model | Architecture | Training Data | | | Generation Modality | | |
|---|---|---|---|---|---|---|---|
| | | DNA | RNA | Protein | DNA | RNA | Protein |
| CD-GPT | Transformer decoder | ✓ | ✓ | ✓ | ✓ | ✓ | ✓ |
| Evo | StripedHyena | ✓ | × | × | ✓ | ✓ | ✓* |
| DNAGPT [10] | Transformer decoder | ✓ | × | × | ✓ | × | × |
| ProtGPT2 [11] | Transformer decoder | × | × | ✓ | × | × | ✓ |
| ProGen [12] | Transformer decoder | × | × | ✓ | × | × | ✓ |
| RNAGEN [13] | WGAN-GP | × | ✓ | × | × | ✓ | × |
| GenerRNA [14] | Transformer decoder | × | ✓ | × | × | ✓ | × |

[2] Zhihan Zhou, Yanrong Ji, Weijian Li, Pratik Dutta, Ramana Davuluri, and Han Liu. Dnabert-2: Efficient foundation model and benchmark for multi-species genome. *arXiv preprint arXiv:2306.15006*, 2023.

[12] Ali Madani, Bryan McCann, Nikhil Naik, Nitish Shirish Keskar, Namrata Anand, Raphael R Eguchi, Po-Ssu Huang, and Richard Socher. Progen: Language modeling for protein generation. *arXiv preprint arXiv:2004.03497*, 2020.

[13] Furkan Ozden, Sina Barazandeh, Dogus Akboga, Sobhan Shokoueian Tabrizi, Urartu Ozgur Safak Seker, and A Ercument Cicek. Rnagen: A generative adversarial network-based model to generate synthetic rna sequences to target proteins. *bioRxiv*, pages 2023–07, 2023.

[14] Yichong Zhao, Kenta Oono, Hiroki Takizawa, and Masaaki Kotera. Generrna: A generative pre-trained language model for de novo rna design. *Plos one*, 19(10):e0310814, 2024.
